# The Impact of Science Beliefs, Teaching Efficacy, and Incentives on Preschool Science Teaching: A Moderated-Mediation Model

**DOI:** 10.1101/2025.11.26.690867

**Authors:** Yanhui Feng, Luqing Wang

## Abstract

Preschool teachers’ ability and willingness to adopt scientific practice teaching are affected by internal and external factors. A question therefore arises: Do organizational incentives, as an external environmental factor (extrinsic motivation), strengthen or weaken the effect of science beliefs (internal factor) on preschool teachers’ scientific practice teaching? We conducted a study with 1,331 teachers from kindergartens in China using voluntary, online, and anonymous questionnaires, implemented through the questionnaire Star application. Evidence shows that science beliefs have a significant positive effect on scientific practice teaching, with teaching efficacy mediating this relationship. Organizational incentives have a significant negative moderating effect on the relationship between science beliefs and scientific practice teaching and teaching efficacy. In this way, organizational incentives and scientific beliefs have a substitutional effect on scientific practice teaching and teaching efficacy. Therefore, preschools with lower incentive levels should aim to implement effective organizational incentives and target teachers’ beliefs. Those with higher incentive levels should ensure their incentives are effective and enhance teachers’ science beliefs to further strengthen teachers’ scientific practice teaching.

## Introduction

Science education is crucial for enhancing national technological competitiveness, cultivating innovative talents, and improving public scientific literacy. The 2011 American publication *A Framework for K-12 Science Education: Practices, Crosscutting Concepts, and Core Ideas* and the 2013 *Next Generation Science Standards* emphasized the importance of enabling students to understand and master core scientific concepts through practical engagement [1]. In recent years, based on a comprehensive understanding of international trends in science education development, China has attached great importance to science education, proposing the working principle of “student-orientation, tailored teaching, and promoting inquiry-based science education.” China’s *Guidelines for the Learning and Development of Children Aged 3–6* (hereinafter referred to as the “Guidelines”) points out, in their educational recommendations, that teachers should support and encourage young children to actively engage both their hands and minds in the process of inquiry to seek answers or solve problems [2]. Children should not only participate in problem-solving, planning, decision-making, and group discussions during exploratory activities but also take part in evaluation activities. The Guidelines emphasize the importance of children’s active involvement in scientific practices.

Teachers are the frontline actors in implementing the Chinese government’s science education policies and are the fundamental force in building a high-quality science education system. Teachers’ proficiency in terms of scientific practice teaching can improve students’ science learning processes and outcomes. This teaching method helps cultivate their critical thinking, ability to conduct collaborative inquiries, scientific attitudes and concepts, and other core components of scientific literacy [3].

Scientific practice teaching is a method through which teachers guide, inspire, and motivate students to ask questions based on learning situations, requiring them to independently explore and solve problems. It emphasizes allowing students to solve real-world problems in real contexts, raising their awareness of how to think like scientists and practice like engineers.

Although science teaching in China effectively imparts scientific knowledge, it does not promote students’ understanding of the nature of science [4]. In China, many educators still adhere to a somewhat traditional view of science instruction, often equating it with the delivery of popular science content. Prominent issues in teaching, such as emphasizing instruction over experience, conclusions over processes, and verification over inquiry, continue to persist. This approach may not fully align with contemporary educational practices or global expectations.

Therefore, teachers’ capacity and willingness to implement scientific practice teaching are crucial; this represents a significant innovative practice that poses considerable challenges for educators accustomed to traditional didactic instruction. Although existing studies have identified individual factors influencing teaching innovation—such as motivation, teaching efficacy, relevant skills, and experience—as well as organizational factors including the work environment, organizational climate, benefits, and team operations, the interactions among these diverse internal and external factors are poorly understood. Self-determination theory (SDT), from an organismic-dialectical perspective, describes how external environments facilitate the internalization of both intrinsic and extrinsic motivation, revealing effective pathways for influencing individual motivation and transcending the limitations of motivation theories such as goal-setting theory, which overemphasize the relationship between goal setting and external incentives [5].

According to SDT, external environmental factors can alter individuals’ perceived loci of causality; they may either undermine or enhance intrinsic motivation. The shift from traditional knowledge-transmission approaches to scientific practice teaching is influenced not only by intrinsic motivations such as interest and beliefs but also by extrinsic factors including material rewards, career advancement, and supervisory policies. Grounded in SDT, we developed a framework that elucidates the interactive effects of organizational incentives and scientific beliefs on early childhood teachers’ implementation of science practice instruction. The results can provide valuable insights for improving science education for preschool children in China.

### Science beliefs and scientific practice teaching

Beliefs cognitively structure experience and are often condensed and integrated into schemas or concepts. They define what people consider to be true and guide individual behavior. Teachers’ beliefs are the intellectual pillars of their professional lives; they comprise the professional creeds teachers adhere to, the core elements of their teaching culture, the invisible guides of their behavior, and the drivers of their internal development [6]. Indeed, teachers’ beliefs influence teaching-related concepts, the formulation of teaching goals, the organization of teaching content, and the selection of teaching methods [7]. Teachers are the driving force of curriculum implementation and the key to the success or failure of reforms. Their understanding of and beliefs about the curriculum and children’s development directly affect their work.

Teachers’ science beliefs are composed of their representations and knowledge of the nature, organization, sources, and standards of proof of scientific knowledge; their mastery of the methods of scientific practice; and their beliefs about the meaning and importance of science teaching. Teachers’ beliefs about cognitive styles and learning beliefs significantly affect teaching in the context of disciplines [8]. We thus hypothesize:

Hypothesis H1: Science beliefs significantly positively predict preschool teachers’ scientific practice teaching.

### The mediating role of science teaching efficacy

Science teaching efficacy refers to preschool teachers’ confidence in their ability to organize and implement science teaching. If a person believes that certain behaviors will produce positive results, they will develop a high sense of self-efficacy and, thus, perform the behavior effectively. Science teachers’ subject knowledge, professional skills, and teaching efficacy affect their scientific practice teaching activities. Enhancing their science beliefs will likely improve their self-efficacy and confidence, prompting them to adopt more effective teaching methods and overcome potential teaching obstacles. The higher the science teaching efficacy of preschool teachers, the more frequently they will organize science teaching activities. Hence, science teaching efficacy is likely to improve the development of preschool teachers’ science teaching behaviors and inquiry-based learning [9]. Notably, science beliefs primarily affect teachers’ teaching attitudes; thus, their effect on science teaching behaviors and scientific practice teaching may be indirect. We thus hypothesize:

Hypothesis H2: Science teaching efficacy mediates the relationship between science beliefs and scientific practice teaching.

### The moderating role of organizational incentives

Teachers’ teaching is shaped by a combination of internal and external dynamics. Internal dynamics include teachers’ own ideals, beliefs, will, goals, and values, whereas external dynamics include cultural, institutional, and material factors, such as social and cultural norms, education systems, and families [10]. In teachers’ scientific practice teaching, internal and external dynamics are indispensable, interconnected, and mutually influential elements of motivation. Science beliefs and efficacy are the internal drivers of teachers’ scientific practice teaching; external dynamics include drivers such as professional training and performance evaluation systems. External dynamics may have a positive or negative effect on scientific practice teaching. Indeed, the extrinsic motivation derived from professional teacher training programs accelerates the improvement of teachers’ professional levels but exposes problems, such as low teacher enthusiasm and poor autonomy [11]. Nevertheless, national- and local-level training is time-consuming and labor-intensive and may not significantly improve teachers’ problem-solving abilities, as some programs have encountered resistance from teachers [12]. In other words, issues pertaining to teachers’ professional development are critical.

In reforming preschool science education, organizational incentives embody top-down requirements, which may involve assigning tasks, regulations, and instructions, making teachers feel less respected, recognized, or cared for. In such a situation, the performance pressure and external control factors related to strong restrictive rewards or punishments can increase teachers’ work pressure and generate emotional fatigue, leading teachers (even those with strong science beliefs and a reasonable understanding of scientific knowledge) to increasingly adopt unchallenging traditional teaching methods, reducing the positive effect of science beliefs on scientific practice teaching. Their confidence in their ability to teach science may be undermined, thereby weakening the positive effect of science beliefs on teaching efficacy. We thus hypothesize:

Hypothesis H3: Organizational incentives negatively moderate the positive relationship between science beliefs and a) teaching efficacy and

1. b) scientific practice teaching.

In summary, this study posits that teaching efficacy and organizational incentives form a moderated mediation model linking science beliefs and scientific practice teaching (Figure 1).

**Fig 1.** **Theoretical framework.**

Based on this in-depth exploration of the relationship between science beliefs and preschool teachers’ scientific practice teaching, we aim to provide theoretical guidance for improving the development of science teaching abilities among China’s preschool teachers.

## Methods

### Participants

This project employed a stratified sampling method, encompassing various regions, levels, professional experiences, and educational experiences. The questionnaire was applied for at the local education authority and then distributed via managers of various units, with teachers filling it out anonymously. All participantsprovided written informed consent via the questionnaire’s preface.This study involved original data collection between July 3rd to July 11th, 2024, via the anonymous online platform Wenjuanxing .A total of 1,454 preschool teachers participated in the study.After excluding 123 invalid responses (based on a reverse-scored attention check question), 1,331 valid questionnaires were obtained (valid recovery rate = 91.5%). Among the participants, 744 were from urban preschools, 371 from township preschools, and 214 from rural preschools; 344 were from first-class provincial preschools, 246 from first-class municipal model preschools, 440 from first-class municipal preschools, and 23 from second-class municipal preschools. In terms of professional experience, 586 teachers had a working experience of 1–5 years, 410 had 6–10 years of experience, 169 had 11–15 years of experience, and 166 had over 15 years of experience. Moreover, 12 had a high school or vocational high school education, 355 had an associate degree, 951 had a bachelor’s degree, and 13 had completed postgraduate education.

### Instruments

#### Science Cognitive Belief Scale

We used the Science Cognitive Belief Scale developed by Niu to measure science beliefs [13]. This scale comprises 12 items covering three dimensions: certainty of scientific knowledge (four items); source of scientific knowledge (five items); and children’s ability to learn science (three items), for example, “Every young child possesses the ability to make inferences and explanations about natural phenomena.” Responses were scored on a 5-point Likert scale ranging from 1 (completely disagree) to 5 (completely agree). The higher the total score, the higher the degree of science beliefs. In this study, Cronbach’s α coefficient was 0.707.

#### Science teaching efficacy scale

The Teaching and Learning International Survey Teacher Questionnaire was used to measure teaching efficacy. It includes 12 items across three dimensions (with four items per dimension): classroom management efficacy, teaching efficacy, and student engagement efficacy. Each dimension contains four items; for example, “I am capable of designing effective scientific inquiry practices for young children.” Answers were rated on a 5-point Likert scale ranging from 1 (completely disagree) to 5 (completely agree). The higher the total score, the higher the teacher’s sense of teaching efficacy. The relevant Cronbach’s α was 0.924.

#### Scientific practice teaching scale

Nine items related to science teaching from the PISA 2015 questionnaire were used to describe the level of scientific practice teaching; for example, “I enable children to design their own scientific experiments or investigation plans.” The items were scored on a 5-point Likert scale ranging from 1 (completely disagree) to 5 (completely agree). Higher total scores indicate higher levels of scientific practice teaching. The resultant Cronbach’s α was 0.963.

#### Organizational incentives scale

Items from the scale developed by Liu and Shi were used to describe the level of organizational incentives [14]. This scale includes four items, such as “Kindergartens have incentive policies to encourage teachers to engage in scientific practice teaching, including promotion systems or salary systems.” The items were scored on a 5-point Likert scale ranging from 1 (completely disagree) to 5 (completely agree). The higher the total score, the higher the incentives for scientific practice teaching in preschools. Cronbach’s α was 0.949 for this scale.

### Data analysis

SPSS^®^ 27.0 was used for descriptive and correlation analysis as well as reliability analysis. PROCESS 4.2 structural equation modeling and regression models were used to test the mediating and moderating effects of teaching efficacy and organizational incentives between scientific beliefs and scientific practice teaching. PROCESS 4.2 was used to test the moderated mediation effect of teaching efficacy and organizational incentives between science beliefs and scientific practice teaching.

## Results

### Common method bias test

A single-factor Harman analysis extracted five common factors with eigenvalues greater than 1, with the first component explaining 22.16% of the total variance—far less than the recommended threshold of 40% [15]. Thus, no issues with common method bias were observed.

### Descriptive statistics and correlation analysis

Pearson’s correlation analysis was conducted on science beliefs, teaching efficacy, organizational incentives, and scientific practice teaching. We found significant positive correlations among all the variables (Table 1).

**Table 1.** Descriptive and correlation analysis (*n* = 1331).

|  | Mean | Standard<br>Deviation | 1 | 2 | 3 | 4 |
| --- | --- | --- | --- | --- | --- | --- |
| Science Beliefs | 3.636 | 0.573 | 1 |  |  |  |
| Science Teaching Efficacy | 3.735 | 0.710 | 0.677** | 1 |  |  |
| Organizational Incentives | 3.883 | 0.824 | 0.598** | 0.725** | 1 |  |
| Scientific Practice<br>Teaching | 3.751 | 0.749 | 0.621** | 0.798** | 0.792** | 1 |

\*\*\* $p < 0.001$ , \*\* $p < 0.01$ , \* $p < 0.05$ .

### Moderated mediation test

First, the interaction terms of the variables were standardized and centered. PROCESS 4.2 (Models 4 and 8) was used to test the moderated mediation model while controlling for teachers’ teaching experience, title and preschool level. We conducted bootstrapping with 5,000 resamples and estimated 95% confidence intervals. Following established procedures for testing moderated mediation effects [16, 17], this study was conducted as follows.

The first step involved testing the simple mediation effect. To explore the underlying mechanism of the significant positive effect of science beliefs on scientific practice teaching, teaching efficacy was further introduced as a mediating variable. It was incorporated into the structural equation model, and Model 4 in PROCESS 4.2 was used to test its mediating effect.

According to Table 2, the upper and lower limits of the bootstrap 95% confidence interval for the mediating effect of teaching efficacy did not include 0, indicating that science beliefs affect scientific practice teaching directly and indirectly. The direct effect accounts for 23.76% of the effect, showing that most of the influence (76.24%) was conveyed through the mediating variable (teaching efficacy).

**Table 2.** Decomposition of total, direct, and mediation effects.

|  | Effect Size | SE | LLCI | ULCI | Effect Size |
| --- | --- | --- | --- | --- | --- |
| Total Effect | 0.621 | 0.022 | 0.579 | 0.663 |  |
| Direct Effect | 0.148 | 0.022 | 0.104 | 0.191 | 23.76% |
| Mediation Effect | 0.474 | 0.026 | 0.424 | 0.525 | 76.24% |

LLCI: lower limit of confidence interval; ULCI: upper limit of confidence interval; S.E.: standard error

Table 3 indicates that the independent variable, science beliefs, has a highly significant positive effect on the mediating variable, teaching efficacy (β = 0.677, *t* = 33.515, *p* < 0.001). Teaching efficacy had a highly significant positive effect on the dependent variable, scientific practice teaching (β = 0.699, *t* = 31.654, *p* < 0.001). The direct effect of science beliefs on scientific practice teaching was also significant (β = 0.148, *t* = 6.679, *p* < 0.001). Supporting the results above, teaching efficacy significantly mediated the influence of science beliefs on scientific practice teaching; the indirect path via teaching efficacy comprised the largest part of the total effect.

**Table 3.** Regression estimation results: The mediating effect of science teaching efficacy.

| Regression Equation | Model Fit Indices | Effect Size and Significance | 95% Confidence |
| --- | --- | --- | --- |

|  |  |  |  |  |  |  |  | Interval |  |
| --- | --- | --- | --- | --- | --- | --- | --- | --- | --- |
| Dependent Variable | Independent Variable | R | R<sup>2</sup> | F | $\beta$ | t | P | LLCI | ULCI |
| Science Teaching Efficacy | Teacher Experience | 0.678 | 0.459 | 281.720*** | 0.025 | 1.259 | 0.208 | -0.014 | 0.064 |
|  | Teacher Title |  |  |  | 0.009 | 0.791 | 0.429 | -0.013 | 0.030 |
|  | Kindergarten Level |  |  |  | 0.014 | 0.956 | 0.339 | -0.015 | 0.042 |
|  | Science Cognitive Belief |  |  |  | 0.677 | 33.515** | 0 | 0.638 | 0.717 |
| Scientific Practice Teaching | Teacher Experience | 0.807 | 0.650 | 493.091*** | -0.022 | -1.346 | 0.179 | -0.053 | 0.010 |
|  | Teacher Title |  |  |  | -0.006 | -0.729 | 0.466 | -0.024 | 0.011 |
|  | Kindergarten Level |  |  |  | -0.016 | -1.352 | 0.177 | -0.038 | 0.007 |
|  | Science Cognitive Belief |  |  |  | 0.148 | 6.679*** | 0 | 0.104 | 0.191 |
|  | Science Teaching Efficacy |  |  |  | 0.699 | 31.654** | 0 | 0.656 | 0.743 |

LLCI: lower limit of confidence interval; ULCI: upper limit of confidence interval. \*\*\* $p < 0.001$ , \*\* $p < 0.01$ , \* $p < 0.05$ .

In the second step, we tested for moderated mediation effects. Based on the hypotheses, we introduced the moderating variable, organizational incentives, into the structural equation model. Using Model 8 in PROCESS 4.2 software, we tested whether organizational incentives moderated the associations between the independent variable (science beliefs) and the mediating variable (teaching efficacy) and between science beliefs and the dependent variable, scientific practice teaching.

As Table 4 illustrates, organizational incentives had a significant predictive effect on both scientific practice teaching (β = 0.430, *t* = 20.353, *p* < 0.001) and teaching efficacy (β = 0.492, *t* = 22.908, *p* < 0.001). The interaction term of organizational incentives and science beliefs significantly negatively predicted scientific practice teaching (β = −0.027, *t* = −2.559, *p* < 0.05) and teaching efficacy (β = −0.030, *t* = −2.368, *p* < 0.05).

**Table 4.** Moderated mediation testing.

| Regression Equation |  | Model Fit Indices |  |  | Effect Size and Significance |  |  | 95% Confidence Interval |  |
| --- | --- | --- | --- | --- | --- | --- | --- | --- | --- |
| Dependent Variable | Independent Variable | R | R<sup>2</sup> | F | $\beta$ | t | P | LLCI | ULCI |
| Science Teaching Efficacy | Teacher Experience | 0.788 | 0.621 | 361.442*** | 0.031 | 1.841 | 0.066 | -0.002 | 0.064 |
|  | Teacher Title |  |  |  | -0.005 | -0.521 | 0.602 | -0.023 | 0.013 |
|  | Kindergarten Level |  |  |  | 0.015 | 1.265 | 0.206 | -0.008 | 0.039 |
|  | Science Belief |  |  |  | 0.377 | 17.822*** | 0 | 0.336 | 0.419 |
|  | Organizational Incentives |  |  |  | 0.492 | 22.908*** | 0 | 0.450 | 0.534 |
|  | Science Belief<br>*Organizational<br>Incentives |  |  |  | -0.030 | -<br>2.368<br>* | 0.018 | -0.054 | -0.005 |
| Scientific<br>Practice<br>Teaching | Teacher Experience | 0.8<br>59 | 0.737 | 530.16<br>5*** | -0.010 | -<br>0.725 | 0.469 | -0.038 | 0.017 |
|  | Teacher Title |  |  |  | -0.016 | -<br>2.082<br>* | 0.038 | -0.031 | -0.001 |
|  | Kindergarten Level |  |  |  | -0.011 | -<br>1.071 | 0.285 | -0.031 | 0.009 |
|  | Science Belief |  |  |  | 0.062 | 3.166<br>** | 0.002 | 0.024 | 0.101 |
|  | Science Teaching<br>Efficacy |  |  |  | 0.438 | 19.12<br>5*** | 0 | 0.393 | 0.483 |
|  | Organizational<br>Incentives |  |  |  | 0.430 | 20.35<br>3*** | 0 | 0.389 | 0.471 |
|  | Science Belief<br>*Organizational<br>Incentives |  |  |  | -0.027 | -<br>2.559<br>* | 0.011 | -0.047 | -0.006 |

LLCI: lower limit of confidence interval; ULCI: upper limit of confidence interval. \*\*\*p <
0.001, \*\*p < 0.01, \*p < 0.05.

Table 5 shows that when organizational incentives were low (M-1SD), teaching efficacy significantly mediated the link between science beliefs and scientific practice teaching, with an indirect effect of 0.178. The mediating effect was also significant when organizational incentives were high (M+1SD), with an indirect effect of 0.152. That is, the strength of the indirect effect of science beliefs on scientific practice teaching through teaching efficacy differs according to the level of organizational incentives; thus, organizational incentives significantly moderate the mediating effect of teaching efficacy between science beliefs and scientific practice teaching.

**Table 5.** Organizational incentives’ moderation of the mediating effect of teaching efficacy.

| Organizational Incentives | Mediation Effect Size | Boot SE | Boot LLCI | Boot ULCI |
| --- | --- | --- | --- | --- |
| M+1SD | 0.152 | 0.020 | 0.116 | 0.193 |
| M | 0.165 | 0.019 | 0.129 | 0.204 |
| M-1SD | 0.178 | 0.021 | 0.138 | 0.221 |

LLCI: lower limit of confidence interval; ULCI: upper limit of confidence interval; SE: standard error.

As discussed above, the interaction between science beliefs and organizational incentives significantly predicted teaching efficacy and scientific practice teaching. Simple slope analysis further demonstrated the predictive role of science beliefs on teaching efficacy and scientific practice teaching at different levels of organizational incentives.

As evident in Figure 2, the influence coefficient of teachers’ science beliefs on teaching efficacy in preschools with high organizational incentives was significantly smaller than that in kindergartens with low organizational incentives. In preschools with relatively low organizational incentives, teachers’ science beliefs more strongly affected their teaching efficacy. Thus, organizational incentives and science beliefs had a substitutional effect on teaching efficacy.

**Fig 2.** **Moderating role of organizational incentives between science beliefs and teaching efficacy.**

As reflected in Figure 3, the influence coefficient of teachers’ science beliefs on scientific practice teaching in preschools with high organizational incentives was significantly smaller than in preschools with low organizational incentives. Hence, in preschools with relatively low organizational incentives, teachers’ science beliefs more strongly affected their scientific practice teaching. Thus, as with teaching efficacy, organizational incentives and science beliefs had a substitutional effect on scientific practice teaching.

**Fig 3.** **Moderating role of organizational incentives between science beliefs and science teaching practice.**

### Conclusions of the results

To briefly recapitulate the main findings: 1) The science beliefs of preschool teachers significantly predicted their scientific practice teaching behaviors; 2) teaching efficacy acted as a mediator between science beliefs and scientific practice teaching behaviors; 3) organizational incentives negatively moderated the impact of science beliefs on a) preschool teachers’ scientific practice teaching and b) teaching efficacy. That is, science beliefs and organizational incentives had a substitutional impact on scientific practice teaching and teaching efficacy.

## Discussion

### Relationship between science beliefs and scientific practice teaching

This study found a significant positive correlation and predictive effect between preschool teachers’ science beliefs and scientific practice teaching, verifying H1 and aligning with the perspective of cognitive-behavioral theory. Brown and Cooney suggested that beliefs define tendencies for action and are the main determinants of behavior [18]. Therefore, teachers’ beliefs can effectively predict their teaching practices [19]. Furthermore, Anderson found that teachers’ beliefs about the purpose of science education can strongly influence their practices [8].

If teachers acknowledge the fundamental trait of scientific knowledge, that is, that such knowledge could be falsified through further testing, they will position themselves and their students as “participants,” actively guiding children to explore the origins and implications of knowledge. This will enable them to implement scientific practice teaching. Educational beliefs are formed during teachers’ pre-service education and continue developing through their professional lives. Therefore, enhancing preschool teachers’ science beliefs by various training programs to improve their level of science teaching is essential.

### The mediating mechanism of teaching efficacy

We found a significant correlation between teachers’ science beliefs, teaching efficacy, and scientific practice teaching; teaching efficacy played a mediating role between science beliefs and scientific practice teaching, confirming H2 and aligning with the motivational theory perspectives. Deehan et al. found a positive correlation between the enhancement of science teaching efficacy gained through university science education courses and the scientific practice teaching adopted by participants in their post-service teaching phase [20]. Enochs and Riggs showed that pre-service elementary teachers exhibit low science teaching efficacy owing to a lack of understanding of scientific concepts and negative science learning experiences during their middle and high school years [21]. Teachers with high teaching efficacy are not only proactive in their work but are also more willing to adopt innovative teaching methods [22].

Additionally, teachers’ sense of teaching efficacy profoundly affects their expectations and guidance of students [23]. Teachers with low science teaching efficacy may fear teaching science and avoid hands-on teaching methods [24]. Conversely, teachers with high teaching efficacy are more likely to implement advanced teaching strategies in the classroom that encourage student autonomy, cater to personalized learning needs, and promote active engagement in classroom tasks [25]. Therefore, helping preschool teachers with low science teaching efficacy to develop their science beliefs can enhance their sense of teaching efficacy and indirectly improve their scientific practice teaching.

### The negative moderating mechanism of organizational incentives

Organizational incentives significantly negatively moderated the first half of the mediating path (i.e., the predictive effect of preschool teachers’ science beliefs on teaching efficacy). Organizational incentives also significantly negatively moderated the direct path between preschool teachers’ science beliefs and scientific practice teaching, confirming H3.

Teachers’ intrinsic motivation for self-development may be constrained by external factors, hindering their professional development. External constraints entail a tension between intrinsic and extrinsic motivations [10]. When imposed by top–down teaching reforms, scientific practice teaching may burden teachers with cognitive and emotional pressure—for example, by forcing them to change established teaching behaviors, adding new tasks to an already heavy teaching workload, and requiring them to convince both themselves and parents and colleagues that new approaches will have positive effects [26]. If preschools adopt an overly prescriptive orientation and fail to recognize that preschool teachers are individuals, this may have a negative effect [27]. If schools focus only on one dimension of science teaching and impose tasks solely from the perspective of pursuing academic achievement without valuing teachers’ individual differences, then the incentive policies (promotion and compensation systems) and training programs provided by preschools will undermine teaching efficacy and scientific practice teaching. Such a prescriptive approach may weaken the positive effect of forms of intrinsic motivation, such as science beliefs and teaching efficacy. Thus, it is crucial to incorporate teachers’ varied professional aspirations, interests, and directions while also inquiring about their views on the optimal components of science education.

Interpersonal incentives significantly enhance teachers’ work engagement, whereas organizational management has not shown a significant effect [28]. Notably, a former director of Sony Corporation, Amami Shiro, proposed that the pursuit of quantifiable performance goals led to the disappearance of passion, love of challenges, and team spirit within Sony, causing the company to lose its status as an innovator and become a laggard [29]. In addition to the in-depth advancement of teaching reform, teaching management in schools has become increasingly scientific, standardized, and regulated. Against such a background, detailed teaching management is increasingly favored. It is important to bear in mind that detailed teaching management can improve the quality of school education: If every aspect of teaching were formally standardized and arranged and teaching is micro-managed, it would negate the individuation and uniqueness of teachers’ teaching and may even result in a decline of educational quality [30].

## Conclusion

### Recommendation 1: Implement organizational incentives that intrinsically motivate preschool teachers’ scientific practice teaching

SDT suggests that psychological needs are the core components connecting the external environment with individual motivation and behavior. When the external environment supports the fulfillment of personal psychological needs, it enhances intrinsic motivation and facilitates the internalization of extrinsic motivation [31]. Accordingly, preschools should align tangible rewards, commands, and directives that intrinsically motivate teachers’ scientific practice teaching. Understanding their specific psychological needs, especially the need for autonomy in scientific practice teaching, is a prerequisite. This approach will leverage the subjective initiative of preschool teachers, returning more decision-making power and authority to them in the field of science education. It would allow them to use their educational creativity and teaching wisdom to make strategic choices, encouraging the proactive and innovative execution of decisions rather than passive adaptation. This transformation from quantity to quality of incentives can enhance teachers’ intrinsic science beliefs and empower their scientific practice teaching.

### Recommendation 2: Integrate theory and practice to strengthen the intrinsic motivation for scientific practice teaching

Beliefs are the driving force behind teachers’ professional development, and a lack of internal motivation can lead to resistance [32]. Science beliefs are formed during teachers’ pre-service education and continue to develop throughout their educational practice. Accordingly, teacher education schools should offer science education courses that integrate theory with practice, allowing teachers in training to experience inquiry-based learning through both theoretical and field-based activities. This will enhance the science beliefs of preschool teachers during their pre-service training and stimulate their intrinsic motivation for scientific practice teaching.

Preschool teachers should be encouraged to recognize the importance of their science beliefs. Through active practice and regular self-reflection, they can continually enrich and transform their science beliefs and discard those that do not align with contemporary scientific education philosophies. By mastering the methods of scientific practice teaching and accumulating experience and skills in science education, they can improve their scientific education capabilities and adaptability, enhancing their confidence in scientific practice teaching and their sense of teaching efficacy. This, in turn, will promote the development and increase the frequency of their scientific practice teaching, advancing the quality of science education in preschools.

### Limitations and prospects

Although the hypotheses of this study were largely substantiated, we must acknowledge certain limitations. First, the questionnaire data were self-reports from preschool teachers; despite conducting tests to mitigate this limitation, the effect of common method variance cannot be entirely ruled out. Future research may consider using observer reports to reduce biases.

Second, during model testing, other variables that may significantly affect preschool teachers’ science teaching practice, such as region, policy, knowledge background, and education level, were not controlled for; this may have influenced the results. Subsequent research should consider the impact of more control variables on the model hypotheses.

Third, the causal relationships between the dimensions of the variables were not considered, as the study design was cross-sectional. Adopting a diverse range of methods, such as latent variable modeling, tracking, and experiments to verify the hypotheses, may help overcome this limitation.

## Acknowledgments

We sincerely thank all kindergarten teachers and principals who participated in this research for their valuable support.

